# Channelrhodopsin variants engage distinct patterns of network activity

**DOI:** 10.1101/275644

**Authors:** Na Young Jun, Jessica A. Cardin

## Abstract

Channelrhodopsins (ChRs) are light-gated ion channels that enable cell type-specific activation of neurons or neural circuits. Channelrhodopsin-2 has been widely used as a tool to probe circuit function *in vitro* and *in vivo*. Several recently developed ChR variants are characterized by faster kinetics and reduced desensitization. However, little is known about how their varying properties may regulate their interaction with local network dynamics. We compared ChR-evoked patterns of multi-unit activity and local field potentials in primary visual cortex of mice expressing three ChR variants with distinct temporal profiles: Chronos, Chrimson, and ChR2. We assessed overall activation of by measuring the amplitude and temporal progression of evoked spiking. Using gamma-range (30-80Hz) LFP power as an assay for local network engagement, we examined the recruitment of cortical network activity by each tool. All ChR variants caused light-evoked increases in firing *in vivo*, but each demonstrated different temporal patterning of evoked activity. In addition, the three ChRs had distinct effects on cortical gamma-band activity. Our findings suggest that variations in the kinetics of optogenetic tools can substantially affect their efficacy in neural networks *in vivo*, as well as the manner in which their activation engages circuit resonance.

## 1. Introduction

The advent of easily accessible optogenetic tools for manipulating neural activity has substantially altered experimental neuroscience. The current optogenetics toolkit for neuroscience comprises a large number of Channelrhodopsins (ChRs), Halorhodopsins, and Archaerhodopsins that enable activation and suppression of neural activity with millisecond-timescale precision. Within the Channelrhodopsin family, many variants have now been made with altered activation spectra, photocycle kinetics, and ion selectivity. The first tool to be widely used in neuroscientific approaches, Channelrhodopsin-2 (ChR2), is a nonspecific cation channel with sensitivity to blue light. ChR2 conferred the ability to evoke action potentials with high precision and reliability across a wide range of cell types.^1-3^ However, the utility of this tool has been somewhat limited by its relatively long offset kinetics and fairly rapid inactivation of photocurrents in response to sustained strong light stimulation. ^1,4-6^ In addition, most naturally occurring Channelrhodopsins are sensitive to blue-green light, presenting a challenge to the use of multiple tools for simultaneous optogenetic control of distinct neural populations. A significant effort in the field has therefore been made to develop Channelrhodopsin variants with faster on- and offset temporal kinetics, less desensitization over time, and red-shifted wavelength sensitivity.

Previous work has suggested that the Channelrhodopsins are highly effective tools for probing the cellular interactions underlying intrinsically generated patterns of brain activity. Stimulation of Parvalbumin (PV)-expressing interneurons in the cortex via ChR2 evokes gamma oscillations, entrains the firing of excitatory pyramidal neurons, and regulates sensory responses.^2,7^ Similarly, ChR2 stimulation of PV^+^ GABAergic long-range projection neurons in the basal forebrain generates gamma-range oscillations in frontal cortex circuits.^8^ Recent work further suggests that ChR2 activation of Somatostatin-expressing interneurons, which synapse on both PV^+^ cells and excitatory neurons, evokes cortical oscillations in a low gamma range.^9^ Sustained depolarization of excitatory sensory cortical neurons via ChR2 activation likewise evokes gamma oscillations, likely by engaging reciprocal interactions with local GABAergic interneurons. ^10^ In comparison, activation of pyramidal neurons in mouse motor cortex via ChRGR, another ChR variant, evokes activity in a broad range of lower-band frequencies.^11^ High-fidelity spiking recruited by Chronos, oChiEF, and ReaChR has been used *in vitro* and *in vivo* in visual cortex ^12-14^ and the auditory midbrain ^15,16^, but the impact of such stimulation on the surrounding network remains unclear.

Despite the substantial increase in available ChR variants with diverse kinetic and spectral properties, it remains unclear how these properties interact with endogenous temporal patterns of neural circuit activity like gamma oscillations *in vivo*. Furthermore, the properties of optogenetic tools are typically tested using short pulses (1 to 100ms) under quiet conditions *in vitro*, but these tools are widely used for sustained neural activation (100s of ms to s) under active network conditions *in vivo*. Here we tested the impact of optogenetic tool properties on evoked activity patterns in the intact brain. We took advantage of the well-characterized gamma oscillation rhythm in mouse primary visual cortex *in vivo* ^10,17,18^ as a metric for optogenetic recruitment of local network activity. Using optogenetic activation of excitatory pyramidal cells as a paradigm to evoke both spiking and cortical gamma oscillations, we compared three Channelrhodopsins with robust photocurrents but distinct kinetic profiles: Chronos, with high-speed on and off kinetics ^19^; ChR2, with fast on but relatively slow off kinetics ^1^; and Chrimson ^19^, with slow on and off kinetics. We found that these tools, although expressed in the same cell type in the same brain region and effective at eliciting action potentials, evoked distinct patterns of activity and had different effects on gamma rhythms. Together, our data suggest that the kinetic properties of engineered opsin tools affect optogenetic interactions with local circuit activity and should be a key factor in experimental design.

## 2. Methods

### 2.1 Animals

All animal handling and maintenance was performed according to the regulations of the Institutional Animal Care and Use Committee of the Yale University School of Medicine. We used both female and male C57BL/6J mice ranging from 3-5 months old.

### 2.2 Surgical procedures

To express ChR2, Chronos, and Chrimson in pyramidal neurons, we injected AAV5-CAMKII-ChR2-GFP (Addgene # 26969), AAV5-CAMKII-CHRONOS-GFP (Addgene # 58805), or AAV5-CAMKII-CHRIMSON-GFP (Addgene # 62718),respectively, in the cortex of C57BL/6J mice. For the virus injection surgery, 1µl AAV was injected through a small burrhole craniotomy in the skull over the left visual cortex [-3.2mm posterior, -2.5mm lateral, -500µm deep relative to bregma] at a rate of 10µl/min using a glass pipette. Injections were made via beveled glass micropipette at a rate of ∼10 nl/min. After injection, pipettes were left in the brain for ∼5 minutes to prevent backflow. Mice were given four weeks for virus expression prior to experiments.

### 2.3 Electrophysiological recordings

Mice were anesthetized with 0.3-0.5% isoflurane in oxygen and head-fixed by cementing a titanium headpost to the skull with Metabond (Butler Schein). All scalp incisions were infused with lidocaine. A craniotomy was made over primary visual cortex and electrodes were lowered through the dura into the cortex. All extracellular multi-unit and LFP recordings were made with an array of independently moveable tetrodes mounted in an Eckhorn Microdrive (Thomas Recording). Signals were digitized and recorded by a Digital Lynx system (Neuralynx). All data were sampled at 40kHz. All LFP recordings were referenced to the surface of the cortex. LFP data were recorded with open filters and MU data were recorded with filters set at 600-9000Hz.

Optogenetic stimulation was provided via an optical fiber (200um) coupled to a laser (Optoengine) at either 470nm (ChR2 and Chronos stimulation) or 593nm (Chrimson stimulation). In each experiment, the fiber was placed on the surface of the dura over the virus injection site and the tetrodes were placed immediately posterior to the fiber.

During each experiment, a total of 150 laser pulses (470 or 593nm) of 1.5s duration were given at varying light intensities (0.5-10mW/mm^2^) with 10s inter-pulse intervals. Bouts of 30 pulses were separated by 5-minute baseline periods.

### 2.4 Histology

Mice were perfused with 0.1M PBS followed by 4% PFA in 0.1M PBS. After perfusion, brains were postfixed for 8 hours in 4% PFA. Brains were sliced at 40μm on a vibratome (Leica) and mounted on slides with DAPI mounting solution (Vector). Images were taken at 10x on an Olympus microscope and the channels were merged using ImageJ (NIH). Laminar distribution of opsin expression was estimated based on DAPI staining.

### 2.5 Data Analysis

Data were analyzed using custom scripts written in Matlab (The Mathworks) and Igor Pro (Wavemetrics). Spikes were detected from the MU recordings using a threshold of +3 SD above the mean, where both the mean and SD were calculated from 10 seconds of recording preceding any light stimulation. Detected spikes were then used to calculate peristimulus time histogram (PSTH) and raster plots for visualization of optically evoked spiking. All firing rate measurements were normalized to the firing rate during a 10-second period prior to all light stimulations. Paired measurements were then taken for the pre-stimulus baseline period prior to each light pulse and the first 1 second of the light-evoked response to that light pulse. For each light intensity, a two-tailed unpaired t-test was performed on the normalized firing rates in the baseline and evoked conditions to determine the presence or absence of an evoked change in firing rate. Inter-spike Intervals (ISI) were calculated as the time interval between successive spikes and a cumulative distribution of ISIs in the on-pulse and off-pulse periods was calculated for each data set

Spectrograms of LFP activity were obtained using 400ms-long Hann windows sliding by 10ms. Prior to STFT, the mean was subtracted to remove DC bias. Each trial was normalized by dividing by the RMS amplitude of the 1s window preceding onset of the light pulse. Spectrograms were averaged across the LFP responses to 28 pulses of 10mW/mm^2^ of 1.5s duration. Relative power in the frequency band of interest was then calculated per frequency bin, setting the average power in the first 0.5 s in each bin to be 1.To further evaluate the changes in gamma range (30-50Hz) activity evoked by optogenetic stimulation, we calculated the ratio of the power spectral density in this frequency band during baseline and stimulation conditions.

The LFP signals included low-amplitude, additive line noise at 60Hz. The method used by Burns et. al. (2010) was not applicable because (1) the amplitude difference caused by the line noise at 60Hz in the full spectrum was not significant enough, and (2) the 60Hz line noise was wide-band and leaked to the neighboring frequency bins of the spectrogram as well. Instead, we noticed that the line noise caused a constant phase shift at the 60Hz line on the spectrogram. The amplitude and phase of the line noise was estimated from the mean value of the complex spectrogram at 60Hz over all time bins during the 4-second interval, assuming that the true 60Hz signal coming from LFP would not have a significant phase bias over the period. When subtracted the estimated line noise from the 60Hz frequency bin as well as the two neighboring bins, this method effectively eliminated the artifact on the spectrogram coming from the 60Hz line noise.

### 2.6 Statistics

For most comparisons, a two-tailed t-test was used. In cases where nonparametric statistics were appropriate due to non-normal data distributions, a two-tailed Kolmogorov-Smirnov test was used.

## 3. Results

### 3.1 Cell type-specific expression of Channelrhodopsins in mouse visual cortex

To understand the efficacy and utility of recently developed Channelrhodopsin variants with differing kinetic properties, we compared three tools: Channelrhodopsin-2, Chronos, and Chrimson (Fig. 1a). We expressed each tool using an AAV construct, under the control of the excitatory neuron-specific CaMKII promoter, into the visual cortex of wild-type mice. Four weeks after virus injection, each of the three Channelrhodopsins was robustly expressed in a characteristic distribution of excitatory pyramidal neurons in cortical layers 2/3, 5, and 6 (Fig. 1c).^2,20^ In each case, opsin expression was widespread in visual cortex, covering up to a distance of up to about 410 µm from the initial injection site. (Fig.1b)

**Figure 1.**
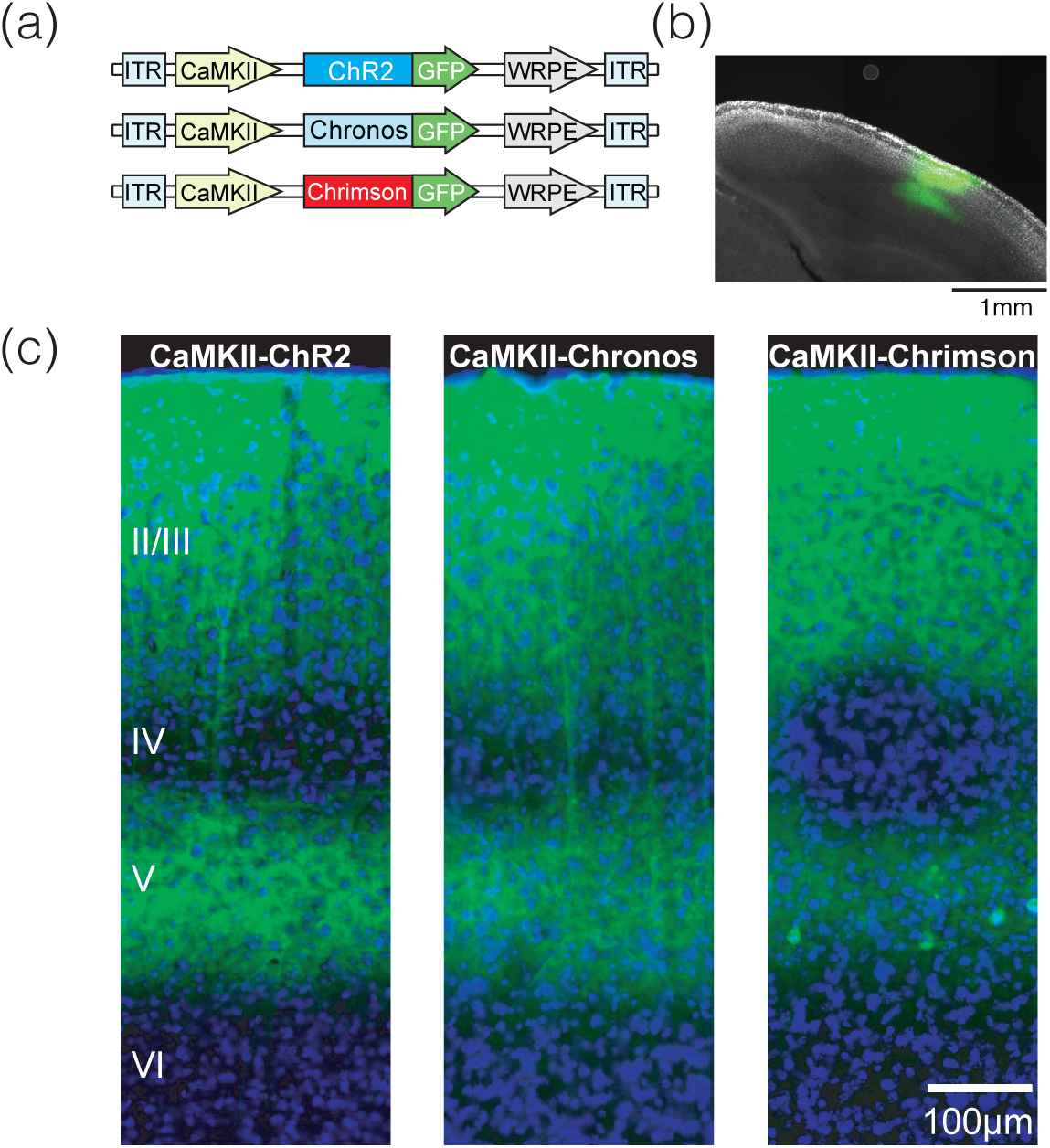
Cell type-specific expression of three Channelrhodopsin variants in excitatory neurons of the mouse visual cortex. **(a)** AAVs carrying three opsins were injected into primary visual cortex of wild-type mice. ChR2, Channelrhodopsin-2; ITR, inverted terminal repeat; WRPE, woodchunk hepatitis B virus post-transcriptional element. **(b)** Example image showing GFP expression (green) around the area of a cortical injection of AAV5 carrying the Chronos construct. Star (⋆) indicates the virus injection site. Magnification: 4x. **(c)** CaMKII-ChR2, CaMKII--Chronos, and CaMKII-Chrimson were robustly expressed in excitatory neurons in cortical layers 2,3, and 5. (Magnification: 10x)

### 3.2 Different Channelrhodopsins evoke distinct cortical activity profiles in vivo

The temporal profile of circuit activity evoked by different opsins may differentially engage network dynamics. To assess the initial and sustained levels of spiking evoked by each opsin, we recorded population multiunit (MU) and local field potential (LFP) activity at multiple cortical sites around each viral injection (ChR2: 11 sites in 3 animals, Chronos: 5 sites in 3 animals, Chrimson: 11 sites in 3 animals). When stimulated with 1.5 seconds of continuous light in an appropriate wavelength (10mW/mm2, Wavelength: 470nm for ChR2 and Chronos, 593nm for Chrimson), all three ChRs evoked sustained firing (Fig. 2a-c). However, each opsin was associated with a distinct temporal profile of spiking.Whereas stimulation of ChR2- (Fig. 2a,d) or Chrimson- (Fig. 2c,f) expressing neurons evoked sustained firing over ∼1-2s, stimulation of Chronos-expressing neurons generated strong initial spiking followed by a decrease towards baseline firing levels (Fig. 2b,e). In contrast, the peak firing evoked by ChR2 and Chronos was rapid and reliable, whereas the peak firing achieved by Chrimson stimulation was delayed and highly variable (Fig. 2d-f inset panels).

**Figure 2.**
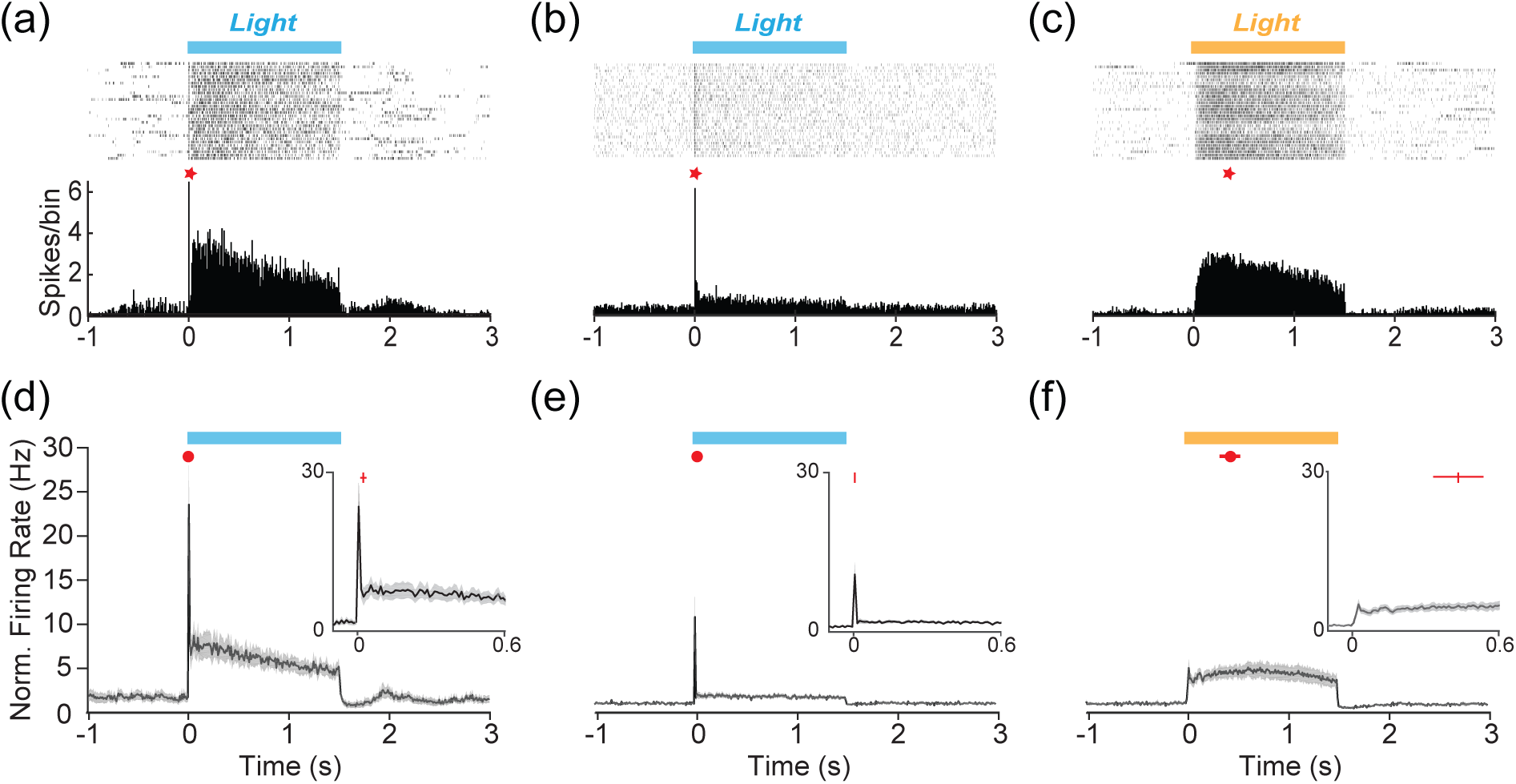
Different Channelrhodopsins evoke cortical activity with distinct temporal profiles *in vivo.* **(a)** Raster plots (upper) and histograms (lower) of example multi-unit spike activity during stimulation of excitatory pyramidal neurons with ChR2. The 1.5s-long interval of light stimulation (10mW/mm^2^) is indicated as shaded box. Star (⋆) indicates the peak firing evoked by the light pulse. **(b)** Same as in (a), for Chronos.**(c)** Same as in (a), for Chrimson. **(d)** Normalized PSTH for all recorded sites in ChR2-expressing mice. Red symbols and lines indicate the mean peak time and s.e.m. of the peak time, respectively. Inset shows the initial period if evoked firing in the first 600ms of light stimulation. **(e)** Same as in (d), for Chronos. **(f)** Same as in (d), for Chrimson.

To quantify these differences in temporal kinetics induced by the three ChR variants, we compared the time between the light pulse onset and the center of the 10 ms interval with the most frequent spikes, averaged over all recording sites and mice for each ChR variant. Chronos showed the shortest peak latency of 0.005 ± 0.01s, whereas Chrimson had a peak latency of 0.43 ± 0.10s, compared to ChR2 0.014 ± 0.01s. (Fig. 2d-f, ChR2: n=3, 11 sites, Chronos: n=3, 5 sites, Chrimson: n=3, 11 sites). The latency to peak was thus shorter for Chronos (p < 0.001) and longer for Chrimson (p < 0.001) compared to ChR2 (Unpaired t-test).

To assay the efficacy of each optogenetic tool in engaging local cortical neurons, we compared the recruitment of spikes in response to a range of illumination intensities. We examined the difference in spike frequency between the baseline and stimulation periods by comparing the distributions of inter-spike intervals (ISIs) evoked by activation of ChR2, Chronos, and Chrimson (Fig. 3). ChR2 (Fig. 3a) and Chrimson (Fig. 3c) both evoked a robust decrease in ISI, consistent with the sustained increase in firing rate, whereas activation of Chronos (Fig 3b) had only a modest effect on the overall ISI distribution. ChR2- (Fig. 3d) and Chrimson- (Fig. 3f) expressing mice showed increasing mean firing rates as the stimulated light intensity increased (linear regression slope: ChR2= 0.41 ± 0.06, p<0.0001, Chrimson= 0.13 ± 0.04, p=0.0025). However, mean firing rates evoked by Chronos activation had a lower rate of increase with light intensity (slope of Chronos = 0.05220 ± 0.01752, p=0.0038). Similar profiles of light intensity-evoked firing responses were observed when analysis was restricted to the initial 100ms of stimulation (linear regresson, slope of ChR2=0.5210 ± 0.06839, p<0.0001, slope of Chrimson= 0.1559 ± 0.03648, p<0.0001, slope of Chronos = 0.06374 ± 0.02281, p=0.0065; Sup Fig. 1).

**Figure 3.**
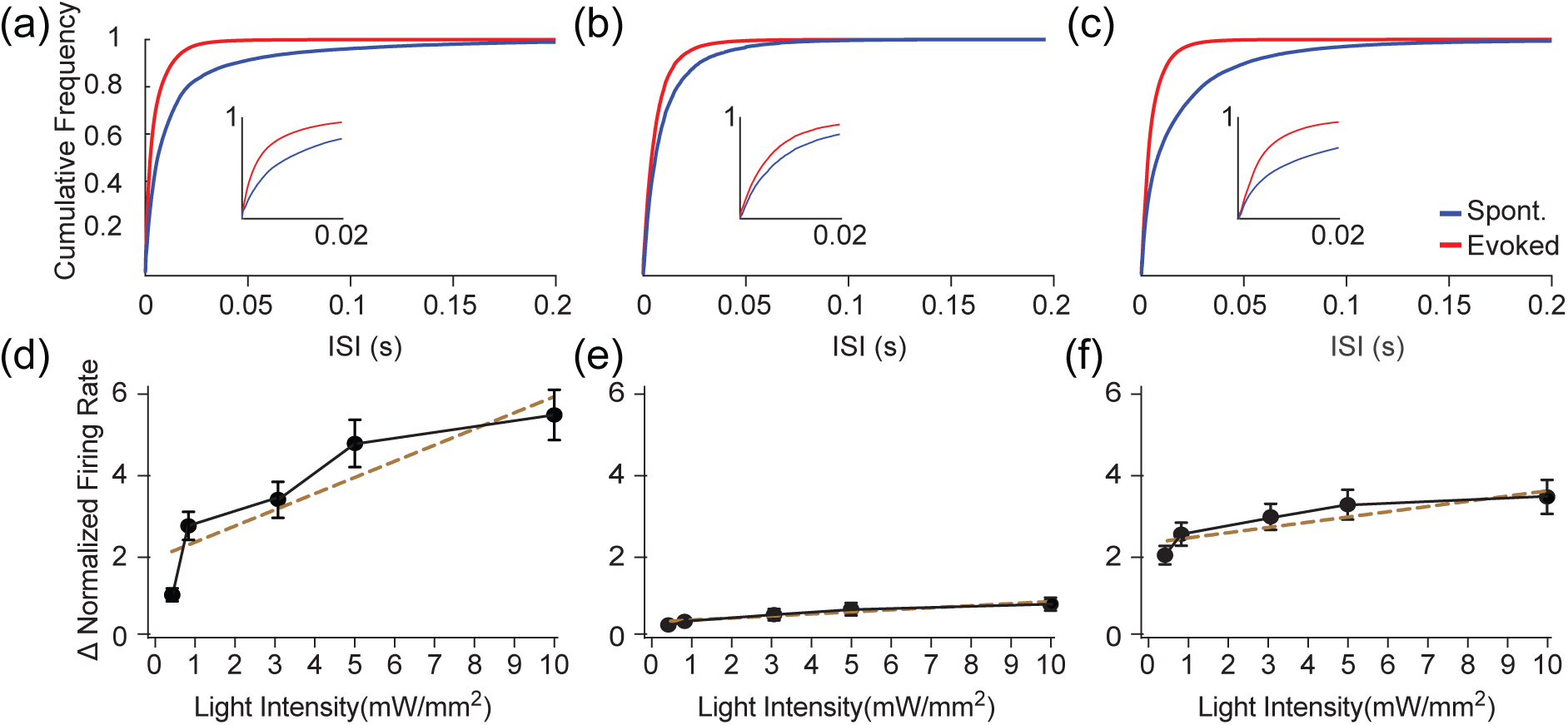
Amplitude and frequency distribution of evoked spike response varies with optogenetic tool. **(a)** ISIs of spontaneous (blue) and evoked (red) MU activity during optogenetic stimulation (10mW/mm^2^) in ChR2-expressing cortex. Inset shows an enlarged plot of the initial 200ms of the evoked spike response. **(b)** Same as in (a), for Chronos. **(c)** Same as in (a), for Chrimson. Error bars denote s.e.m. **(d)** Firing rates evoked by ChR2 stimulation over a range of intensities, normalized to spontaneous firing rates. **(e)** Same as in (d), for Chronos. **(f)** Same as in (d), for Chrimson. Dashed lines indicate linear regression of the data. Error bars denote s.e.m.

### 3.3 Opsin-specific recruitment of cortical gamma rhythms

Previous work has found that ChR2 stimulation of pyramidal neurons engages the cortical gamma rhythm (30-80Hz), an outcome of resonant excitatory-inhibitory circuit interactions^21^, *in vitro* and *in vivo*.^10^ Using evoked gamma power as a measure of network activation, we assayed the efficacy of each optogenetic tool in driving recurrent circuit interactions. Activation of ChR2-expressing excitatory neurons evoked a response in the local field potential (LFP) and a broadband increase in high-frequency activity (Fig. 4a, Sup Fig.2d). Chronos and Chrimson likewise evoked an initial deflection of the LFP signal (Fig. 4b-c; Sup Fig. 2b,c). However, neither Chronos nor Chrimson activation of excitatory neurons evoked the characteristic sustained high-frequency LFP activity observed following ChR2 stimulation of the same population of neurons.

**Figure 4.**
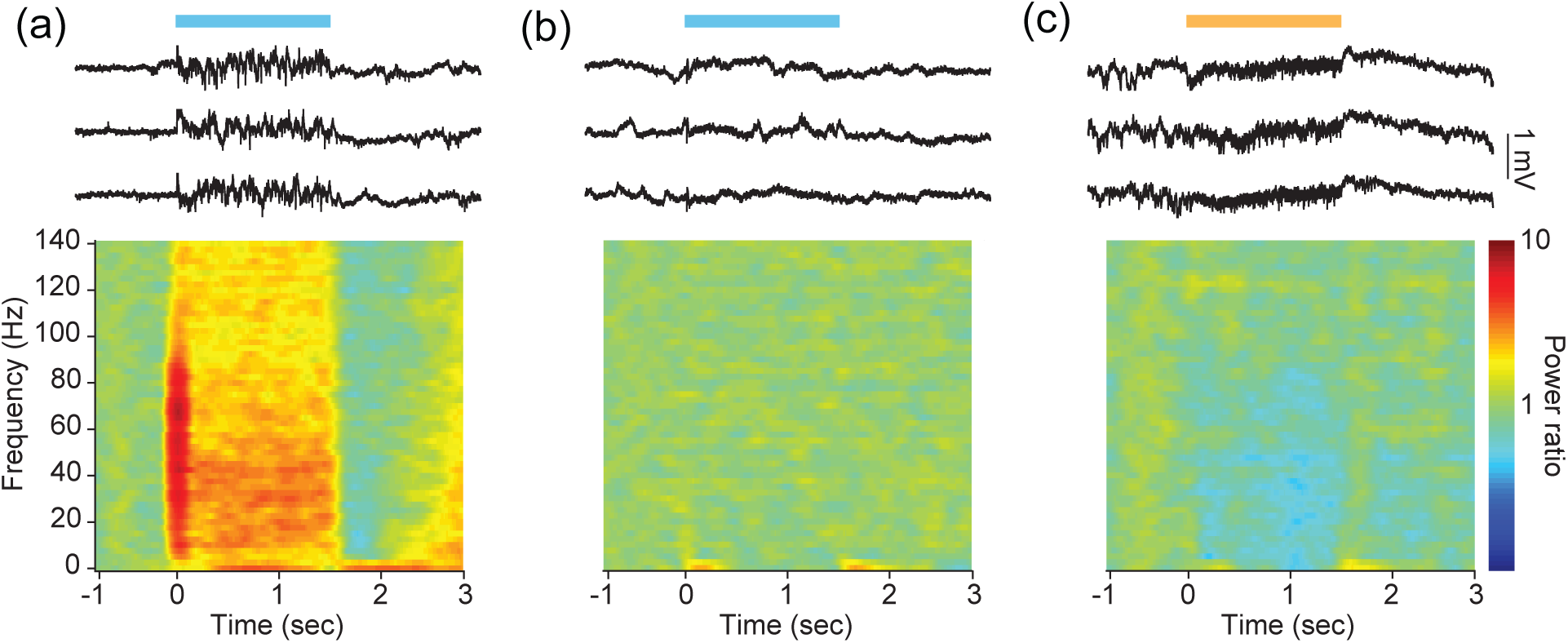
Different Channelrhodopsins evoke distinct cortical activity profiles *in vivo*. **(a)** Example traces of cortical LFP activity in response to 1.5s of 10mW/mm^2^ light stimulation of ChR2-expressing pyramidal neurons (upper). Average changes in spectral power density at this site across stimulation trials (lower). **(b)** Same as in (a), for Chronos. **(c)** Same as in (a), for Chrimson.

In agreement with previous work^10^, we found that the light-activation of ChR2-expressing excitatory neurons amplified local field potential (LFP) power in the gamma range (ChR2: 11 sites in 3 animals; Kolmogorov-Smirnov test, p<0.01; Fig. 5a, d). In contrast, neither Chrimson nor Chronos demonstrated similar engagement of endogenous patterns of cortical network activity during stimulation. Stimulation of excitatory neurons via activation of Chronos increased gamma power at low (< 3mW/mm^2^) but not high (10mW/mm^2^) light intensities (non-parametric test, p=0.6994; 5 sites in 3 animals). Surprisingly, excitatory neuron stimulation via Chrimson significantly suppressed cortical gamma power in a light-intensity dependent manner (p<0.0001; 11 sites in 3 animals). Together, these data suggest that the pattern of activity recruited by stimulation with these three different tools engages distinct modes of endogenous circuit interactions.

**Figure 5.**
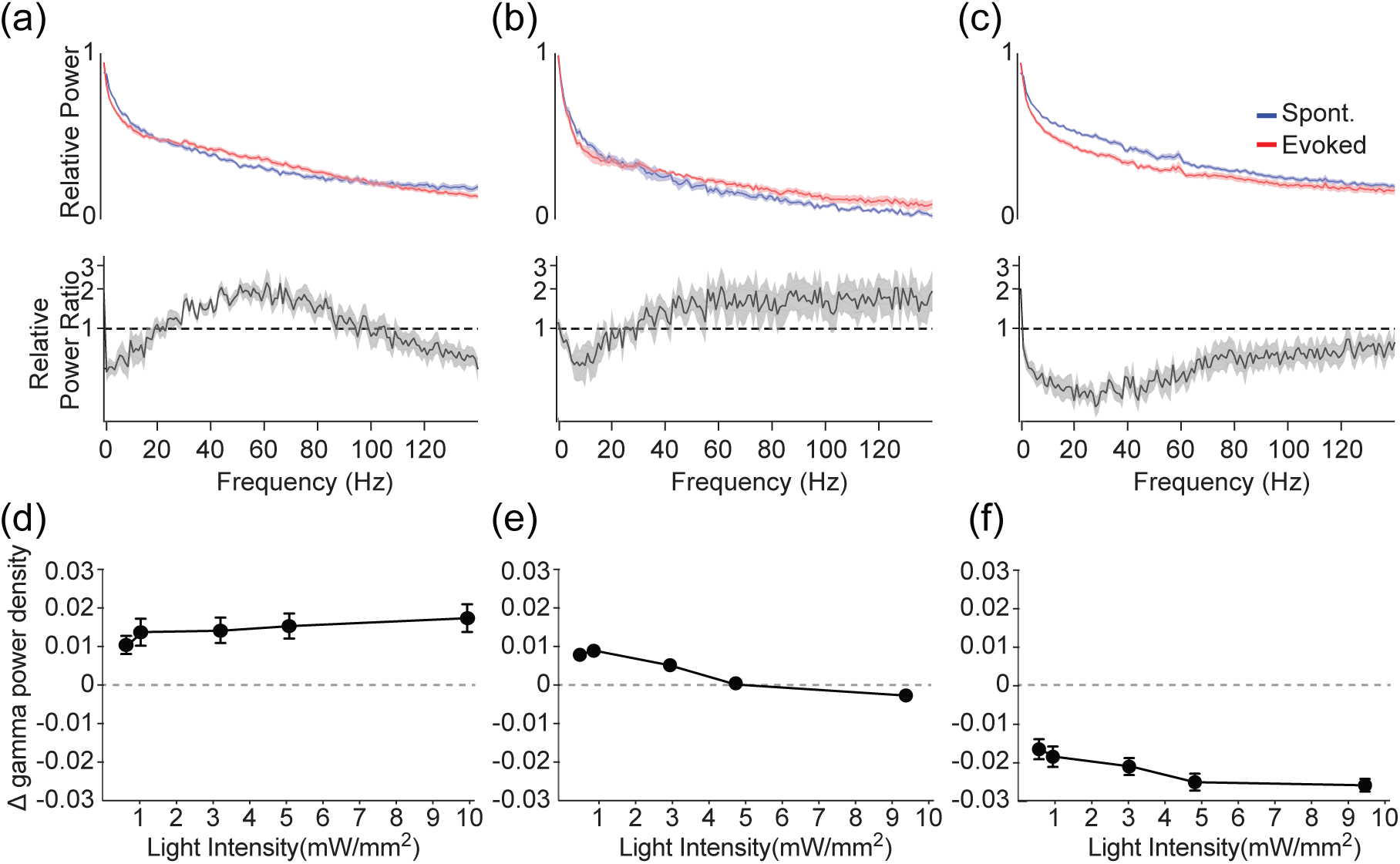
Distinct recruitment of gamma-band activity by different Channelrhodopsins. **(a)** Normalized power spectra (upper) for spontaneous (blue) and evoked (red) cortical LFP activity in response to ChR2 stimulation and the ratio between evoked and spontaneous spectra (lower). **(b)** Same as in (a), for Chronos. **(c)** Same as in (a), for Chrimson. Shaded areas denote ± s.e.m. **(d)** Change in the relative power in the gamma band (30-50Hz) in response to varying light intensities in cortex expressing ChR2. **(e)** Same as in (d), for Chronos. **(f)** Same as in (d), for Chrimson. Error bars denote s.e.m.

## 4. Discussion

Although recent work has resulted in the development of new Channelrhodopsin variants to meet experimental needs, the *in vivo* effects of opsins with distinct properties have not been fully explored. In particular, varying temporal profiles of optogenetically evoked neural activation may substantially affect the manner in which the surrounding neural circuit is engaged. Here we expressed three opsins (ChR2, Chronos, and Chrimson) with different kinetics in excitatory pyramidal neurons in the primary visual cortex. Using a previously validated paradigm for optogenetic recruitment of gamma-range resonance in the local cortical circuit, we compared the temporal envelope of evoked spiking and gamma-range activity across the three opsins. Although all three tools were effective in driving enhanced spiking, the temporal profile of the evoked activity was distinct. In addition, only ChR2 stimulation generated increased cortical gamma activity.

Recently developed Channelrhodopsins vary extensively in their kinetic profiles. ChR2 exhibits a relatively fast onset (τ_on_) but a long offset time (τ_off_), leading to diminished temporal fidelity in spike responses. ^22^ The τ_off_ of several ChRs also slows further upon membrane depolarization.^23^ The cumulative effect of this long τ_off_ is to cause a prolonged depolarization after the evoked action potential, preventing rapid re-hyperpolarization of the membrane and contributing to artificial spike doublets. Prolonged depolarization may also inactivate voltage-gated channels needed for high-frequency spiking. Mutations in ChR2 to accelerate the closure of the pore gave rise to the CheTA variant, which has high temporal fidelity but reduced light sensitivity and charge transfer.^22^ ChRGR, a ChR1 variant, shows rapid τ_on_ and τ_off_, along with reduced desensitization.^11^ In comparison, Chronos and ChIEF exhibit large photocurrents, rapid deactivation, and improved efficacy in eliciting high-fidelity fast spiking.^19,24^

A second series of tools were developed with absorption spectra shifted towards longer wavelengths compatible with 2-photon imaging ^25-27^ and dual-channel optogenetic circuit interrogation.^19^ Initial chimerization between VChR1 and ChR1 gave the red-shifted C1V1^28^, with a red-shifted absorption peak but relatively low photocurrents and very slow kinetics. Further work produced several additional red-shifted tools, including ReaChR^29^, with a peak similar to C1V1 but larger photocurrents, and bReaCHES^30^, a faster variant with larger photocurrents and higher spike fidelity. In comparison, Chrimson exhibits a more red-shifted absorption peak and very large photocurrents,making it highly effective for driving robust neural activity.^19^ Chrimson has substantially slower τ_on_ and τ_off_ properties than ReaChR, bReaCHES, ChR2, or Chronos. Based on their low toxicity, robust expression levels, large peak photocurrents, and distinct kinetic profiles, we selected Chronos and Chrimson for *in vivo* comparison with ChR2.

We found a strong relationship between the properties of the individual opsins and the temporal profile of the spiking they evoked. The two tools with relatively rapid onset kinetics, ChR2 and Chronos, each evoked a precisely timed initial spike event across the neuronal population, followed by sustained spiking at lower firing rates. In contrast, Chrimson, with slow onset kinetics, did not evoke reliable spiking at stimulation onset and gave rise to a much broader temporal distribution of spike frequencies. These results, along with previous findings^6,23^, suggest that the kinetics of the opsins interact meaningfully with intrinsic neuronal membrane properties. Rapid membrane depolarization, like that caused by ChR2 or Chronos activation, contributes to recruitment of voltage-gated channels and enhances the reliability and precision of the initial evoked action potentials in cortical neurons.^31,32^ In comparison, a slow rate of depolarization, like that caused by Chrimson, leads to temporally disbursed spiking. We further found that the kinetics of the opsins shaped the overall temporal envelope of the sustained spiking evoked by long stimulation. Whereas the initial efficiacy of ChR2 and Chronos resulted in an early peak in evoked firing rates within the first 50ms, the spike response to Chrimson stimulation peaked several hundred milliseconds later. However, the sustained firing rates evoked by ChR2 and Chrimson were higher than that evoked by Chronos, suggesting that rapid deactivation of this opsin may reduce overall spike rates.

Gamma oscillations are generated by rhythmic interactions between excitatory and inhibitory neurons.^21^ Activation of excitatory neuron synaptic input to predominantly fast-spiking interneurons causes a highly synchronous and precise spike response in the interneurons, temporarily suppressing excitatory neuron spiking. When excitatory spiking recovers following the inhibitory event, the interneurons are again recruited. This temporally structured, reciprocal interaction between excitation (E) and inhibition (I) leads to a very robust 30-80Hz network oscillation with a ∼25ms period determined by the time course of inhibition. Gamma activity in cortical circuits can be evoked by optogenetically stimulating either the inhibitory interneurons^2,7,9^ or the excitatory neurons.^10,33^ Generation of gamma oscillations by excitatory neuron stimulation likely results from the highly synchronous activation of a large volley of spikes from excitatory neurons, which are highly effective in activating the inhibitory neuron spiking that sets the temporal pattern for resonance in the network.^2,7,21,34^ Several cycles of gamma can be produced by even a single brief stimulation of excitatory pyramidal neurons^7^, but sustained gamma oscillations in active cortical networks *in vivo* may require consistently elevated excitatory spiking.^34,35^ Given these temporal constraints, the different temporal profiles of evoked excitatory neuron spiking evoked by the three ChRs could potentially engage varying network responses.

In good agreement with previous work^10^, we found that ChR2 stimulation of excitatory pyramidal neurons evokes robust cortical gamma activity. In contrast, stimulation of the same neuronal population via Chronos evoked little to no gamma activity, presumably because the initial, highly precise spiking from stimulated cells is not followed by sufficiently elevated excitatory spiking to sustain network oscillations. In comparison, Chrimson might be expected to evoke little gamma because the slow increase in activity precludes an initial burst of spikes. Suprisingly, we found that stimulation via Chrimson also significantly suppressed endogenous gamma. These results suggest that Chrimson’s slow temporal kinetics and late firing peak destabilize the highly precise interplay between E and I cells, increasing the firing rates of excitatory neurons but precluding their entrainment by inhibition.

Overall, we found that differences in the properties of three Channelrhodopsins were associated with distinct profiles of evoked cortical activity. Although this does not represent an exhaustive evaluation of all available opsins, our data suggest that the temporal kinetics of the opsins affect the temporal profile of evoked activity on multiple time scales. Rapid onset kinetics may facilitate the recruitment of highly precise initial spike responses, whereas slow onset kinetics preclude synchronous spiking and result in delayed peak responses. In addition, opsins with distinct kinetics interact differently with endogenous circuit resonance, affecting the sustained patterns of activity evoked in cortical networks over longer time periods. Our findings suggest complex interactions between optogenetic tools and active neuronal networks in the intact brain. The optogenetics toolkit for neuroscience includes an ever-increasing variety of tools with varying properties, and individual tools may be appropriate for different experimental goals.

## Acknowledgments

We would like to thank Dr. Edward Boyden for generously sharing samples of viral vectors for Chronos and Chrimson used in initial pilot experiments. We would also like to thank Mr. Jong Wook Kim for assistance with coding and data analysis. This work was supported by NIH R01 MH102365, NIH R01 EY022951, a Smith Family Award for Excellence in Biomedical Research, a Klingenstein Fellowship Award, an Alfred P. Sloan Fellowship, a NARSAD Young Investigator Award, and a McKnight Fellowship to JAC.

---

**Supplementary Figure 1.**
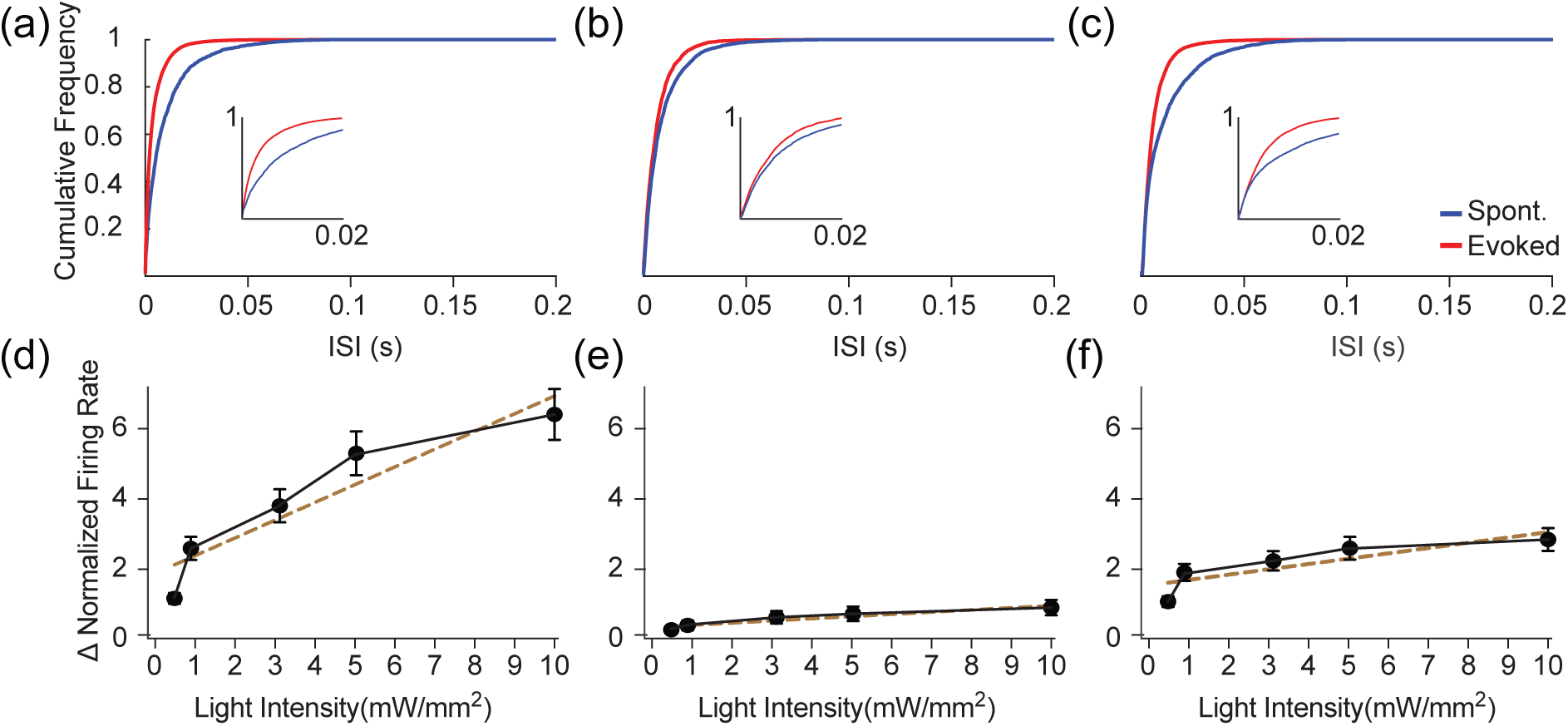
Distinct changes in evoked spike patterns during the first 100ms after light onset. **(a)** ISIs of spontaneous (blue) and evoked (red) MU activity during the initial 100ms of optogenetic stimulation (10mW/mm^2^) in ChR2-expressing cortex. **(b)** Same as in (a), for Chronos. **(c)** Same as in (a), for Chrimson. Error bars denote s.e.m. **(d)** Firing rates evoked by the initial 100ms of ChR2 stimulation over a range of intensities, normalized to spontaneous firing rates. **(e)** Same as in (d), for Chronos. **(f)** Same as in (d), for Chrimson. Dashed lines indicate linear regression of the data. Error bars denote s.e.m.

**Supplementary Figure 2.**
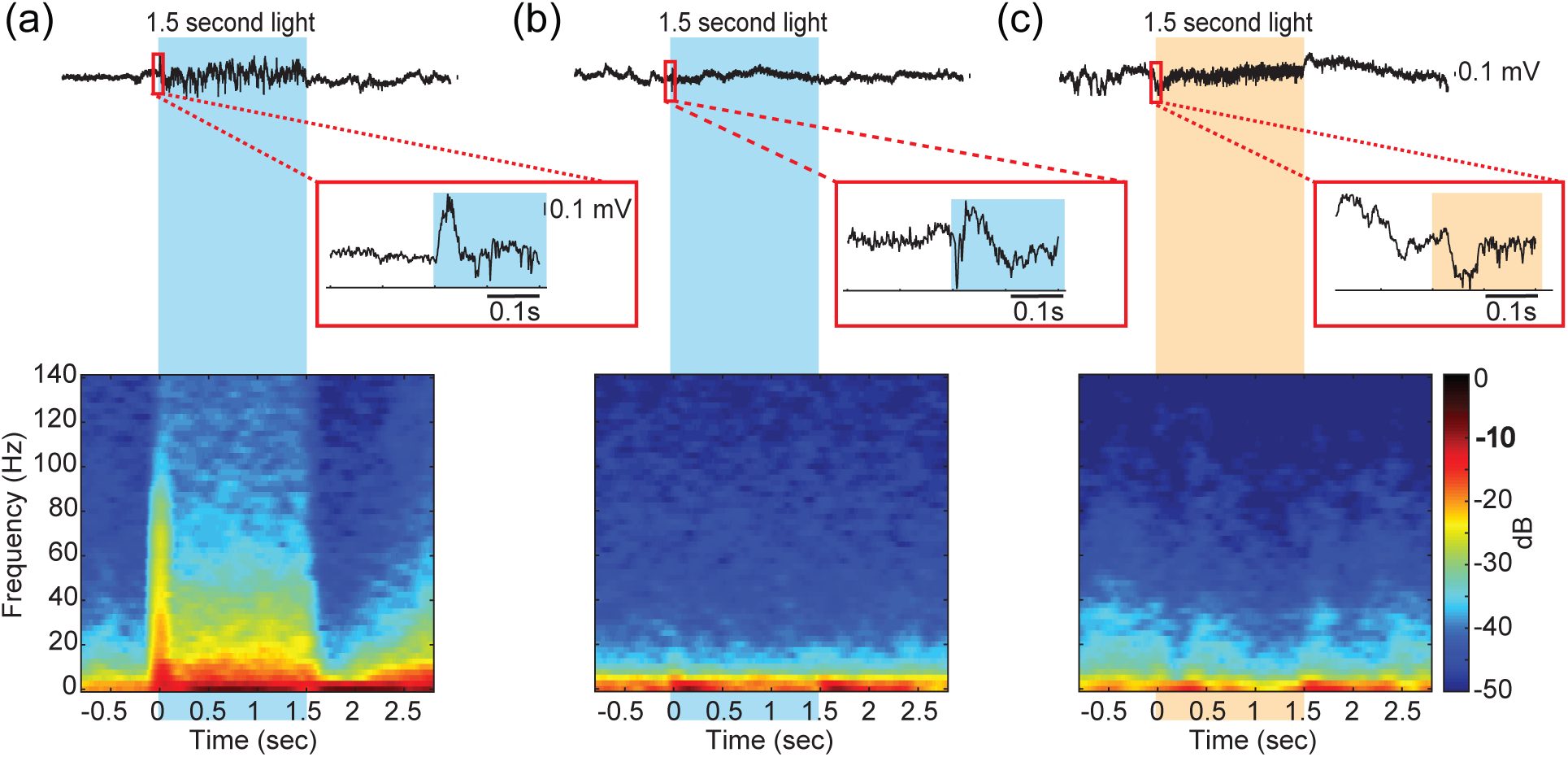
Different Channelrhodopsins evoke distinct cortical activity profiles *in vivo.* **(a)** Example traces of cortical LFP activity in response to 1.5s of 10mW/mm^2^ light stimulation of ChR2-expressing pyramidal neurons (upper). Inset: The LFP traces between 0.2s before and after the light stimulation. **(b)** Same as in (a), for Chronos. **(c)** Same as in (a), for Chrimson. **(d)** LFP spectrogram of the trace in (a), shown as raw spectral power. **(e)** Same as in (d), for Chronos. **(f)** Same as in (d), for Chrimson.

